# Selection on mutators is not frequency-dependent

**DOI:** 10.1101/735100

**Authors:** Yevgeniy Raynes, Daniel M. Weinreich

**Affiliations:** Center for Computational Molecular Biology, and Department of Ecology and Evolutionary Biology, Brown University, Providence, Rhode Island 02906, USA

## Abstract

The evolutionary fate of mutator mutations – i.e., genetic variants that raise the genome-wide mutation rate – in asexual populations is often described as being frequency (or number) dependent. This common intuition suggests that mutators can invade a population by hitchhiking with a sweeping beneficial mutation, but only when sufficiently frequent to produce such a mutation before non-mutators do. Here, we use stochastic, agent-based simulations to show that neither the strength nor the sign of selection on mutators depend on their initial frequency, and while the overall probability of hitchhiking increases predictably with frequency, the per-capita probability of fixation remains unchanged.

## Main text

Mutator alleles have been found at considerable frequencies in populations of infectious and commensal bacteria [1–3], viruses [4], and pathogenic fungi [5–7]. Mutators are also believed to be widespread in in many cancers [8, 9], and have been repeatedly observed to overtake microbial populations during laboratory evolution experiments [10–17]. Yet, unlike directly beneficial mutations that are favored by natural selection because they increase an organism’s reproductive success (i.e. its fitness), mutator mutations generally do not appear to be inherently advantageous [16] (except potentially in some viruses [18, 19]). Instead, mutators experience indirect selection, mediated by persistent statistical associations with fitness-affecting mutations elsewhere in the genome. As a result, mutators may invade an adapting population by hitchhiking [20] with new beneficial mutations even when they have no effect on fitness of their own [21].

Common intuition suggests that whether or not mutators can successfully hitchhike to fixation depends on the initial prevalence of mutator alleles in a population - most commonly referred to as frequency or number dependence [16, 21]. This view holds that to replace the resident non-mutators, mutators must generate a beneficial mutation that escapes genetic drift and sweeps to fixation before their non-mutator competitors do. Accordingly, it has been proposed that mutators may be expected to invade (i.e. are favored by selection) only when already present in sufficient numbers to produce the successful beneficial mutation first, and lose their advantage (i.e. are disfavored by selection) when too rare to do so [16, 21].

This frequency-dependent interpretation of mutator success has found empirical support in studies of mutator hitchhiking in laboratory microbial populations. Most famously, in a series of pioneering experiments, Lin Chao and colleagues showed that mutator strains of the bacterium *E. coli* could supplant otherwise isogenic non-mutator strains by hitchhiking with beneficial mutations when initialized above a critical threshold frequency but would decline when initialized below it [22, 23]. Since then, a similar pattern has been recapitulated in several other studies in *E. coli* and *S. cerevisiae* yeast [24–27].

Here, we demonstrate that, contrary to this widespread intuition, selection on mutators is independent of frequency. To do so, we use individual-based, stochastic computer simulations of asexual populations that mimic microbial evolution experiments under generally-accepted parameter values [28]. Fig. 1A shows mutator frequency dynamics in randomly chosen simulations initialized across four log-orders of starting frequency, *x*_0_, which recapitulate experimental observations of the critical frequency threshold for hitchhiking (i.e. in [22, 23]). Individual realizations started below an apparent threshold frequency all end in mutator loss, while simulations started above end in fixation (Fig. 1A).

**Figure 1:**
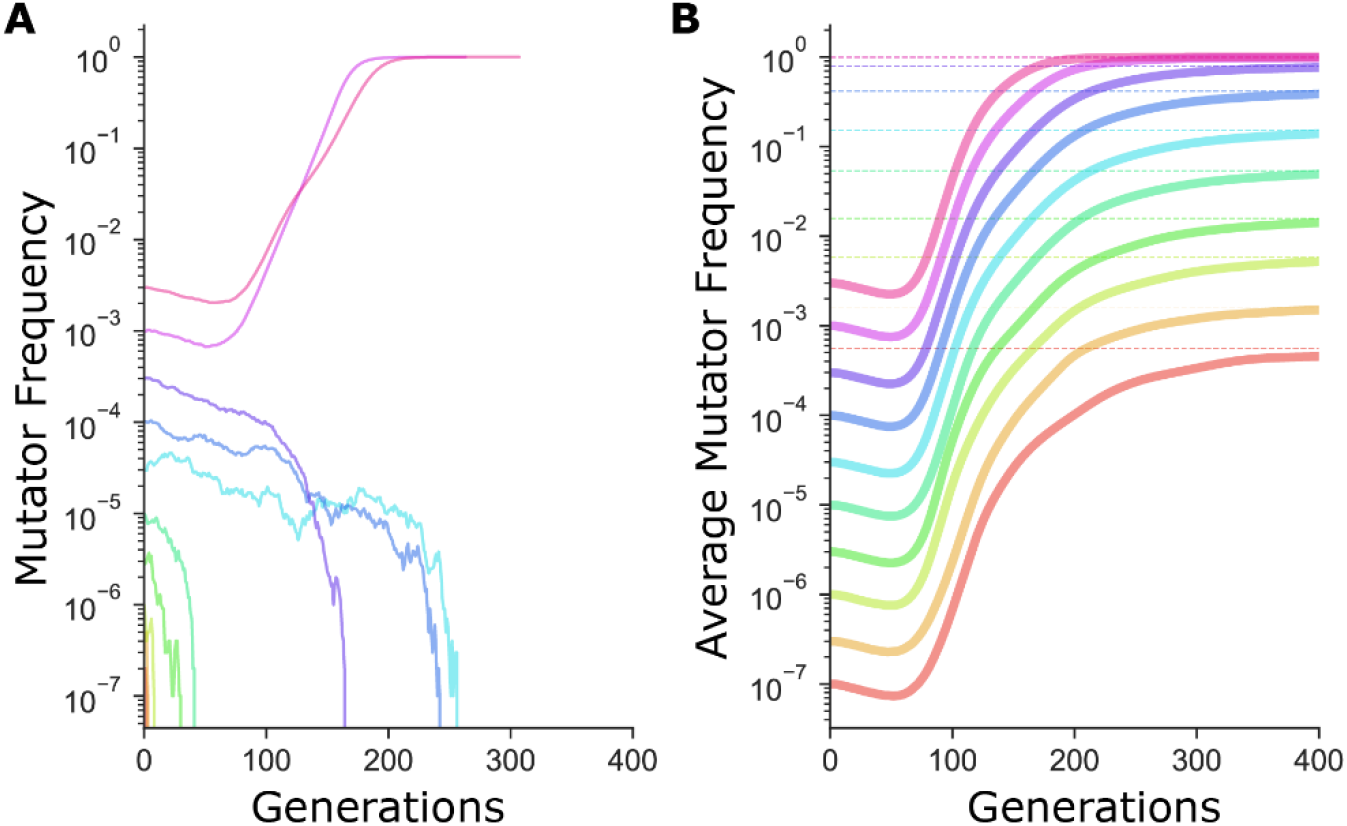
The sharp transition between fixation and loss in mutator dynamics at different starting frequencies is due to limited sampling. **A)** In simulations, mutator trajectories in individual realizations initiated at different starting frequencies recapitulate the experimental observation of the frequency-threshold for mutator hitchhiking. Parameter values used are typical of microbial experimental populations [28]: *N* = 10^7^, *U_del_* = 10^-4^, *U_ben_* = 10^-6^, *s_ben_* = 0.1, *s_del_* = −0.1. Mutators mutate 100× faster than non-mutators. **B)** Average mutator trajectories across realizations do not show evidence of the frequency-threshold. On average, mutators increase in frequency at all *x*_0_, showing that selection favors mutators independent of frequency. Average mutator frequency always eventually reaches the expected *P_fix_*(*x*_0_) (dashed horizontal lines) calculated in Fig 2. Mutator frequencies averaged across 10^6^ simulation runs at *x*_0_ = 10^-7^ and *x*_0_ = 3×10^-7^, and across 10^5^ simulation runs for all other starting frequencies.

Critically, fixation of an allele in a finite population is a probabilistic process influenced both by selection and random genetic drift, and even beneficial mutations will frequently be lost by chance alone. As such, whether an allele is truly favored or disfavored by selection can only be ascertained by evaluating its expected behavior averaged across many replicate, independent realizations. Indeed, if we consider the expected mutator frequency averaged across many replicate simulations, the threshold-frequency effect disappears (Fig. 1B). Instead, the average mutator frequency ultimately rises above the starting frequency at all *x*_0_, suggesting that mutators are, in fact, favored by selection in these populations regardless of starting frequency. [For more on why mutators are favored in large populations such as these see [28], and the significance of the transient decline in frequency seen in Fig. 1B will be explored in a forthcoming publication.]

To confirm that selection on mutators is independent of starting frequency, we measured the fixation probability of a mutator allele, *P_fix_*(*x*_0_), at each initial frequency, *x_0_* simulated in Fig. 1. Given the stochasticity of the fixation process (and following others [28–30]), we compare *P_fix_*(*x*_0_) to the probability of fixation of a neutral allele, given by *x*_0_. If a mutator is favored, we expect it to fare better than neutral (i.e. *P_fix_*(*x*_0_) > *x*_0_), and worse than neutral (i.e. *P_fix_*(*x*_0_) < *x*_0_) if disfavored. As Fig. 2 shows, *P_fix_*(*x*_0_) exceeds the fixation probability of a neutral allele for all *x*_0_, as anticipated in Fig. 1B and confirming that the sign of selection on mutators does not depend on starting frequency.

**Figure 2:**
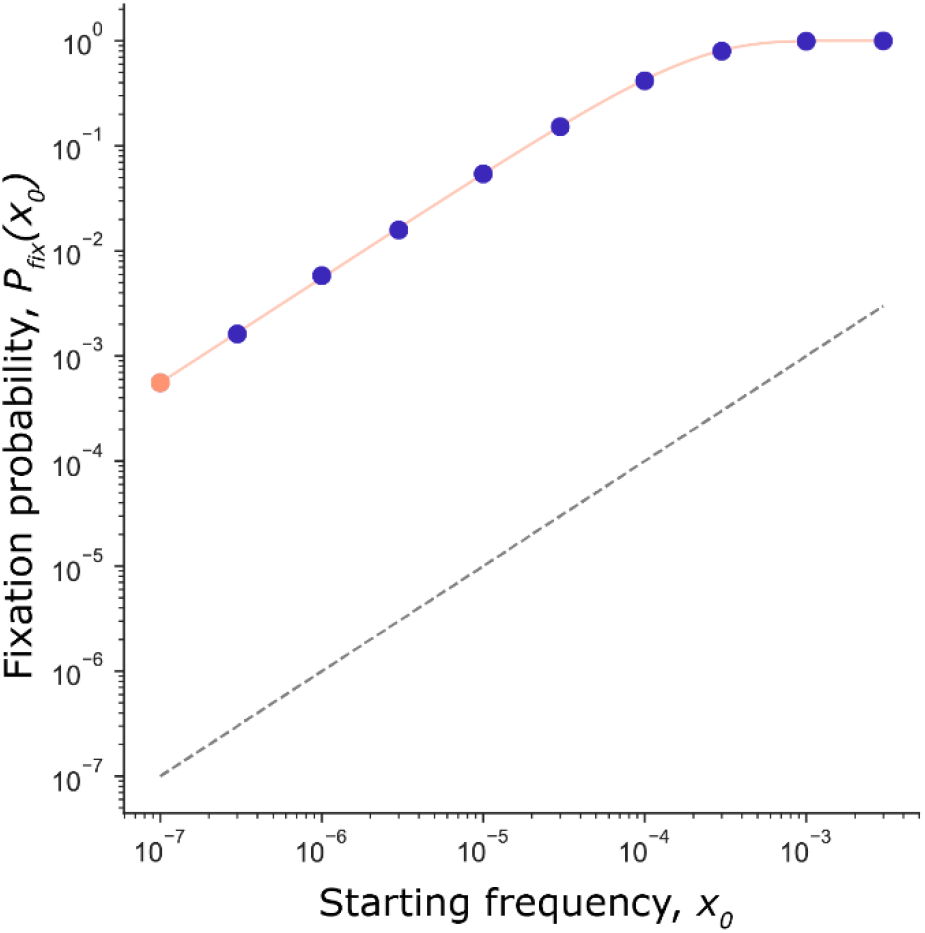
Mutator fixation probability is not frequency-dependent. Fixation probability, *P_fix_*(*x*_0_), of a mutator initiated at frequency *x*_0_ (circles: orange for *x*_0_ = 1/*N*, purple for *x*_0_ > 1/*N*). Data from simulations in Fig. 1. *P_fix_*(*x*_0_) scales with but never crosses the fixation probability of a neutral mutation (*x*_0_; black dashed line). Thus, mutators are favored at all starting frequencies. The expected fixation probability *P_fix_*(*x*_0_) (solid orange line), calculated from the fixation probability of a single mutator, *P_fix_*(*x*_0_ = 1/*N*) = 5.6×10^-4^ (orange point) using Eq. 1 is indistinguishable from the *P_fix_*(*x*_0_) observed in simulations, demonstrating that the per-capita fixation probability at every frequency is independent of *x_0_* and equal to *P_fix_*(*x*_0_ = 1/*N*).

Furthermore, while *P_fix_*(*x*_0_) of a mutator increases with *x*_0_, it does so exactly as expected for a frequency-independent mutation. Under frequency-independent selection *P_fix_*(*x*_0_) is simply the probability that at least one of the *x*_0_*N* alleles reaches fixation (where *N* is the population size). By definition of frequency-independent selection, the per-capita fixation probability is a constant, written, *P_fix_*(*x*_0_ = 1/*N*). Correspondingly, *P_fix_*(*x*_0_) for any *x*_0_ can be calculated as

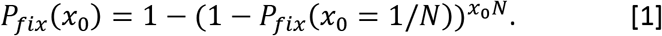

As the orange line in Fig. 2 shows, *P_fix_*(*x*_0_) calculated with Eq. 1 is indistinguishable from *P_fix_*(*x*_0_) observed in simulations, confirming that the per-capita fixation probability is independent of *x*_0_ and equal to *P_fix_*(*x*_0_ = 1/*N*) at any *x*_0_.

Why then do mutators in experimental populations appear destined to go extinct when initially rare (Fig. 1A and [22, 23])? Consider that the per-capita fixation probability of a mutator is relatively low – in our simulations operating under realistic parameter values *P_fix_*(*x*_0_ = 1/*N*) = 5.6×10^-4^. Thus even when mutators are favored, most experimental replicates with rare mutators are expected to end in mutator extinction, and only those started at frequencies higher than roughly 1/[*NP_fix_*(*x*_0_ = 1/*N*)] are expected to end mostly with mutator fixation. Therefore, considering only a single or even a few realizations (as in Fig 1A) would, most likely, result in observing only the most expected outcome for each *x*_0_. Such limited sampling across a broad range of starting frequencies explains the sharp transition between fixation at high frequencies and loss at lower ones even when selection is frequency-independent [see also Discussion in [31]].

In fact, the critical frequency-dependent transition observed in Fig. 1A is not unique to mutators. Recall that *P_fix_*(*x*_0_) of any mutation not under frequency-dependent selection, nevertheless, increases with starting frequency, *x*_0_ (Eq. 1). For example, even for a directly beneficial mutation, the probability of fixation from low frequencies is relatively low (Fig. 3A Inset), Correspondingly, Fig. 3A illustrates how single realizations of the dynamics of a directly beneficial mutation again exhibit a threshold-like switch from fixation to loss. In contrast, expected frequency dynamics averaged across many realizations confirm that beneficial mutations are favored by selection independent of starting frequency (Fig. 3B). Indeed, only for mutations under truly frequency-dependent selection do both the individual realizations (Fig. 3C) and the expected dynamics across many realizations (Fig. 3D) exhibit a frequency-dependent transition.

**Figure 3:**
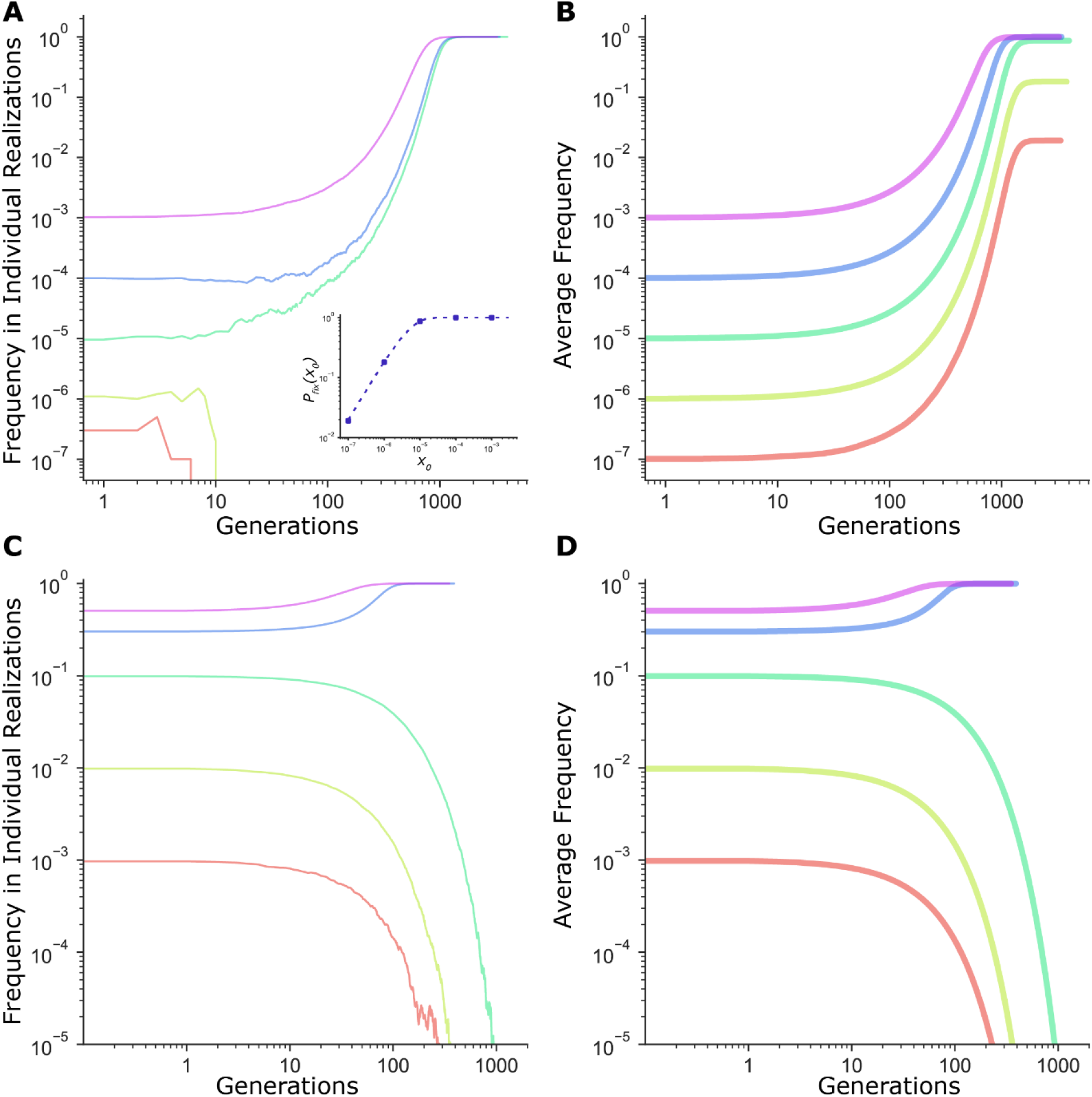
Frequency threshold in dynamics of fitness-affecting mutations. **A)** Individual realizations of a simulation initiated with a directly beneficial mutation of size *s_ben_* = 0.01 at a starting frequency *x*_0_. Population size, *N* = 10^7^. **Inset:** Fixation probability of a beneficial mutation of size *s_ben_* =0.01 at *N* = 10^7^. Dashed line is given by 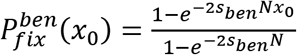 [34], while circles are values of 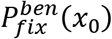 measured in simulations (averaged across 10^5^ runs). **B)** Average frequency trajectories of a beneficial mutation of size *s_ben_* = 0.01 in **A** averaged across all 10^5^ runs of simulation. **C)** Individual realizations of a simulation initiated with a mutation under frequency dependent selection, with the selection coefficient *s*(*x*) = *b* + *mx*, where *x* is the frequency, *b* = −0.02, *m* = 0.1, and *N* = 10^7^. **D)** Average frequency trajectories of the frequency-dependent mutation in **C** averaged across all 10^5^ runs of simulation. All panels are on a log-log scale for clarity.

In summary, our results demonstrate that neither the strength nor the sign of selection on mutators depend on initial frequency or number. Instead, we show that in populations favoring higher mutation rates, mutators consistently fare better than the neutral expectation (Fig. 1 and Fig. 2) regardless of starting frequency. Most importantly, the per-capita probability of fixation remains unchanged with frequency. We conclude that the frequency threshold observed in earlier experiments is, therefore, an artifact of limited experimental sampling rather than a frequency-dependent change in selective effect.

## Methods

Individual-based, stochastic simulations employed here have been previously described [28]. In brief, we consider haploid asexual populations of constant size, *N*, evolving in discrete, non-overlapping generations according to the Wright-Fisher model [33]. Populations are composed of genetic lineages - subpopulations of individuals with the same genotype. A genotype is modeled as an array of 99 fitness-affecting loci and 1 mutation rate modifier locus, which in a mutator state raises the genomic mutation rate *m*-fold. Beneficial mutations at the fitness loci increase fitness by a constant effect *s_ben_*, while deleterious mutations decrease fitness by a constant effect *s_del_*. We assume additive fitness effects and so calculate fitness of a lineage with *x* beneficial and *y* deleterious mutations as *w_xy_* = 1 + *xs_ben_* − *ys_del_*. Simulations start with the mutator allele at a frequency of *x*_0_ and continue until it either fixes or is lost from a population.

Every generation the size of each lineage *i* is randomly sampled from a multinomial distribution with expectation 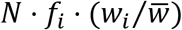, where *f_i_* is the frequency of the lineage in the previous generation, *w_i_* is the lineage’s fitness, and 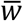 is the average fitness of the population 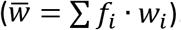. Upon reproduction, each lineage acquires a random number of fitness-affecting mutations M, drawn from a Poisson distribution with mean equal to the product of its size and its total per-individual mutation rate, (*U_ben_* + *U_del_*), where *U_ben_* and *U_del_* are the deleterious and beneficial mutation rates respectively. The number of beneficial and deleterious mutations is then drawn from a binomial distribution with *n*=*M* and *P* = *U_ben_*/(*U_ben_* + *U_del_*) and new mutations are assigned to randomly chosen non-mutated fitness loci.

## Acknowledgements

We thank Paul Sniegowski and Benjamin Galeota-Sprung for comments on the manuscript. Simulations were performed on the computing cluster of the Computer Science Department at Brown University. The work was supported by National Science Foundation Grant DEB-1556300.

